# IS*1216* drives the evolution of pRUM-like multidrug resistance plasmids in *Enterococcus faecium*

**DOI:** 10.1101/2025.07.10.664085

**Authors:** Freya Allen, Ross S. McInnes, Willem van Schaik, Robert A. Moran

## Abstract

pRUM-like plasmids are commonly found in multidrug-resistant *Enterococcus faecium*, but the evolution of these plasmids has not been characterised in detail. When we analysed the genome sequences of two clinical *E. faecium* strains isolated in Birmingham, UK, we found two pRUM-like plasmids, pHHEf1 and pHHEf2. They were approximately 25 kb in size and shared the same 10 kb backbone, but contained starkly different accessory regions that were bounded by and interspersed with the IS*26* family insertion sequence IS*1216*. pHHEf1 contained a complete set of vancomycin resistance genes, while pHHEf2 contained aminoglycoside and erythromycin resistance genes along with an integrated small plasmid, pCOLA. It appeared that IS*1216* had driven the diversification of these accessory regions. We sought to characterise the role of IS*1216* in the broader evolution of pRUM-like plasmids by performing comparative analyses on 152 complete plasmid sequences from five continents. Extensive IS*1216*-mediated variation included backbone deletions, acquisition and loss of 10 different antibiotic resistance genes, and the formation of cointegrates with plasmids of at least 10 different replicon types. Cointegration events have introduced accessory segments with diverse functions, including horizontal transfer determinants and genes for bacteriocin T8. The derivations of these acquired segments highlight the impact of IS*1216* in driving gene exchange between *Enterococcus* and *Staphylococcus* species. We traced the emergence of the pRUM-like lineage to a putative ancestor found in a vancomycin-sensitive ST17 *E. faecium* isolated in 1997. The ancestral plasmid, pCANE, includes the entire pRUM backbone with an additional 44.9 kb in place of the pRUM accessory region. The 44.9 kb segment includes putative conjugation determinants, suggesting that the emergence of the pRUM-like lineage coincided with a loss of transfer functions. We propose an IS*1216* driven model for the evolution of pRUM-like plasmids, which appear to have arisen in *E. faecium* ST17 and contributed to the international success of CC17 as an opportunistic pathogen.

**Impact Statement:** Experimental work has shown that the insertion sequence IS*1216*, found in *Enterococcus*, shares the cointegrate formation properties of the well-characterised IS*26* of Gram-negative bacteria, which is a major contributor to the movement and accumulation of antibiotic resistance genes in important human pathogens. It seems likely that IS*1216* plays a similar role in the emergence and spread of multidrug resistance in *Enterococcus*. However, the actions of IS*1216* in the genus have not been the subject of focused comparative studies. Here, we provide genomic evidence for the crucial role of IS*1216* in the evolution of the pRUM-like multidrug resistance plasmid lineage that has been found in *E. faecium* isolated globally. We have traced the evolution of the pRUM-like lineage to a cryptic ancestral plasmid found in a vancomycin-sensitive human clinical strain isolated in the 1990s.

## Introduction

*Enterococcus faecium* is a commensal resident of the gastrointestinal tracts of humans and animals.^1^ Some *E. faecium* strains can cause opportunistic extra-intestinal infections that range from urinary tract infections to bacteraemia.^2, 3^ These infections are particularly prominent in nosocomial settings, with intensive care units (ICUs) sites of significant concern. *E. faecium* was recognised as a nosocomial pathogen by the 1980s, but its rising prominence in the late 1990s coincided with the acquisition of vancomycin resistance by pathogenic strains. The resistance to last-resort antibiotics such as vancomycin, and the increasing prevalence of multidrug resistance (MDR) in the species severely compromises antimicrobial chemotherapy.^1^ In 2019 over 50,000 deaths were attributed to antibiotic resistant *E. faecium* globally.^4^

*E. faecium* can acquire antibiotic resistance determinants through horizontal gene transfer (HGT). Plasmids are major drivers of HGT, and several plasmid types are known to play an important role in the dissemination of resistance genes in enterococci.^5, 6^ The content and structures of plasmids are shaped by the actions of smaller mobile genetic elements (MGEs), including insertion sequences (IS) and transposons.^7^ As well as carrying accessory genes such as those that confer antibiotic resistance, smaller MGEs can interrupt, delete or invert segments of plasmid backbones. These structural changes can impact both the plasmid and its microbial host by altering functions such as conjugative transfer, and can be utilised in comparative studies to trace the evolution of plasmid lineages.^8, 9^

The plasmid pRUM, first described in 2003, was found in a clinical *E. faecium* that was isolated in Ohio, USA in the 1990s.^10, 11^ The toxin-antitoxin system of pRUM, Axe-Txe, was the focus of this initial study, becoming the first proteic plasmid addiction system to be characterised in Gram-positive bacteria.^11^ In addition to *axe*-*txe*, pRUM contained genes for replication and partitioning, UV resistance, and erythromycin and aminoglycoside resistance.^11^ The pRUM replication initiation gene, *repA*, encodes a RepA_N family replicase, making it part of a group that includes diverse representatives found in various Gram-positive bacteria, including the MDR plasmid pSK41 of *Staphylococcus aureus* and the pheromone responsive plasmid pCF10 of *Enterococcus faecalis*.^12^ pRUM-like plasmids are common in clinical MDR strains of *E. faecium* (clade A1), and have been strongly associated with the problematic clonal complex 17 (CC17).^5, 13–15^ Previous studies have described variation in pRUM-like plasmid content, noting differences in the antibiotic resistance genes (ARGs) carried, the presence of various IS, the structure of the vancomycin resistance transposon Tn*1546*, and in the variable presence of *axe*-*txe* that was first noted in molecular studies.^13–16^

IS*1216* is an 809-bp IS*26* family IS found in enterococci. IS*26* has received significant attention in Gram-negative bacteria, where it is a major driver of resistance gene dissemination and accumulation.^17^ IS*26* exhibits unique properties relative to IS of other families. A particularly important characteristic of IS*26* is its ability to form cointegrates in a targeted and high-efficiency mode.^18^ This conservative mode preferentially targets other copies of IS*26*, which ensures that DNA molecules that contain IS*26* are predisposed to acquiring additional copies, along with any associated accessory genes in the form of translocatable units (TUs).^18^ This phenomenon has been responsible for the accumulation of antibiotic resistance determinants in plasmids and chromosomal islands in various Gram-negative pathogens.^19–21^ Experimental evidence has demonstrated that IS*1216* behaves similarly to IS*26*, and can form cointegrates using both copy-in and conservative modes.^22^ Consistent with this, IS*1216* has also been shown to amplify resistance genes in clinical *E. faecium*, modifying the vancomycin resistance phenotype of its host.^23^

Although IS*1216* has been associated with antimicrobial resistance in *Enterococcus* for some time,^7^ its role in the evolution of plasmids in the genus has not been explored to the extent of IS*26* in Gram-negative bacteria. To that end, we have investigated the pRUM-like lineage of MDR plasmids by examining 152 complete plasmid sequences: two derived from human infections at a hospital in Birmingham, and 150 from various countries that were available from GenBank. By comparing plasmid structures, we have traced the evolution of this lineage, and identified a putative progenitor to it in the complete genome of a clinical *E. faecium* isolate from 1997. This has highlighted the crucial role of IS*1216* in the generation of the pRUM-like lineage, and its ongoing impact on the evolution of these now internationally distributed and clinically relevant plasmids.

## Methods

### Whole genome sequencing and sequence assembly

Illumina sequencing (250 bp paired) was performed on the NovaSeq 6000 platform at MicrobesNG (Birmingham, United Kingdom). For long-read sequencing, DNA was extracted using the Wizard ® Genomic DNA Purification kit (Promega) and quantified using the Qubit Broad Range dsDNA kit (Thermo Fisher). Genomic DNA was prepared with the SQK-LSK109 ligation kit and sequenced on a R9.4.1 flow cell using the MinION platform (Oxford Nanopore Technologies). The complete genome of E1162 was assembled using Unicycler (v. 0.4.8). The sequences of pHHEf1 and pHHEf2 were generated as part of a previous study,^24^ but annotated and submitted to GenBank here.

### Plasmid annotation and sequence comparison

The GenBank non-redundant nucleotide database was screened for pRUM-like plasmids using the sequence of the pRUM *repA* gene, which was obtained from GenBank accession AF507977 (last search 23/11/2023). All complete plasmid sequences that contained identical copies of the pRUM *rep* gene were retained for further study. Host sequence type (ST) was determined using mlst (https://github.com/tseemann/mlst) if its chromosome sequence was available.

In preparation for comparative analyses, all complete plasmid sequences were opened at the same position, which was the first base of *repA*. Plasmids were screened for reference sequences with abricate v0.9.8 (https://github.com/tseemann/abricate), using the ResFinder database for ARGs,^25^ and the PlasmidFinder database for plasmid replicons^26^. IS were identified using ISFinder^27^. Plasmid sequences were compared using standalone BLAST+^28^. Sequence annotations were performed manually and visualised using Gene Construction Kit version 4.5 (Textco Biosoftware Inc.). Once identified, 100 bp junction sequences indicative of specific molecular events were generated by taking 50 bp contiguous sequences from either side of backbone-MGE or MGE-MGE junctions, and used to screen the collection as described previously.^29^

## Results

### Clinical *E. faecium* harbour pRUM-like plasmids carrying different antibiotic resistance genes

Complete pRUM-like plasmid sequences were found in the genomes of two *E. faecium* strains isolated from clinical blood specimens at Birmingham Heartlands Hospital in 2016 and 2017. The plasmids were of similar size but carried different sets of ARGs (Figure 1). pHHEf1, carried by ST262 strain OI5 isolated in 2016, was 25,483 bp and contained a complete set of *vanA*-type vancomycin resistance genes. pHHEf2, carried by ST780 strain OI17 isolated in 2017, was 24,877 bp and contained one erythromycin and three aminoglycoside resistance genes. pHHEF1 and pHHEF2 have the same 10,157 bp backbone that is interrupted by an IS*1216*-bounded accessory region and by a single copy of IS*1678* (Figure 1). The backbone is typical of pRUM-like plasmids and includes replication determinants and the *axe*-*txe* toxin-antitoxin genes,^11^ but does not contain a transfer region or any transfer-associated genes.

**Figure 1:**
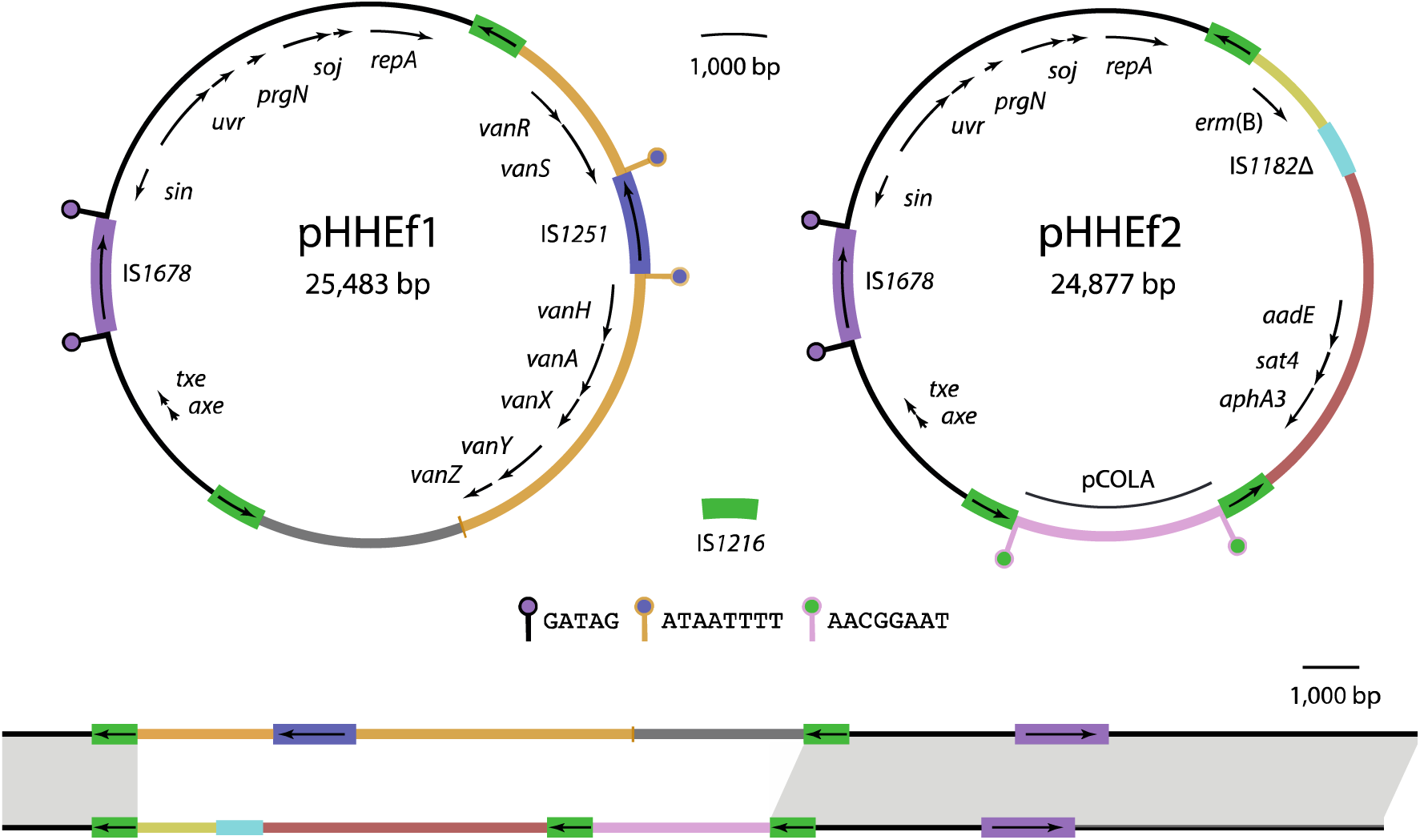
Birmingham pRUM-like plasmids pHHEf1 and pHHEf2. Scaled circular maps of plasmids pHHEf1 and pHHEf2 above a linear alignment of the two. The extents and orientations of genes are indicated by labelled arrows. Segments of different derivations are shown in different colours, with the colours maintained across the circular and linear representations of the plasmids. Insertion sequences are shown as labelled boxes with internal arrows indicating the extents and orientations of their transposase genes. Target site duplications (TSDs) generated by the actions of insertion sequences are shown as ‘lollipop’ shapes with stems coloured according to the DNA segment duplicated and circles according to the element responsible for the duplication. The sequences of TSDs are indicated in the key. The linear representations of the plasmids are opened at the first base of *repA*, and grey shading between the lines that represent the plasmids links shared segments with nucleotide identity >99.9%. Drawn to scale from GenBank accessions PV751131 and PV763145.

The antibiotic resistance regions of pHHEf1 and pHHEf2 are located in the same position within the backbone and bounded by directly oriented copies of IS*1216*. However, the size and content of the resistance regions differ. Excluding the bounding IS*1216*, the resistance region of pHHEf1 is 12,024 bp and comprised largely of a truncated copy of transposon Tn*1546*, which lacks transposition genes but contains a complete set of vancomycin resistance genes, with a copy of IS*1251* inserted between *vanS* and *vanH* (Figure 1) in a configuration that has been described previously.^30^ The resistance region of pHHEf2 is 11,419 bp and contains an additional IS*1216* in the same orientation as the boundary copies. The additional IS*1216* separates 7,403 bp and 3,207 bp segments. The larger of these includes a truncated copy of IS*1182* separating the erythromycin resistance gene *erm*(B) from the aminoglycoside resistance genes *aadE* (also called *ant(6)-Ia*), *sat4*, and *aphA3* (also called *aph(3’)-III*) (Figure 1). The 3,207 bp segment appears to be a small plasmid, which we refer to as pCOLA, that was acquired in an IS*1216*-mediated transposition event. The pCOLA segment is 99.9% identical to the circular 3,199 bp plasmid pVRE32643_5 (GenBank accession OP378676), differing only by a single nucleotide polymorphism and the presence of the 8 bp target site duplication (TSD) AACGGAAT immediately adjacent to the IS*1216* at its boundaries (Figure 1). The TSD sequence lies within the pCOLA *rep* gene, which has been interrupted and separated in the IS*1216*-mediated event that incorporated pCOLA into the pRUM-like cointegrate.

Considered together, the structures of pHHEf1 and pHHEf2 resemble those of characterised IS*26*-associated antibiotic resistance regions in Gram-negative bacteria, including in F-like plasmids in *Escherichia coli* and in chromosomal islands in *Acinetobacter baumannii*.^19, 20^ The loss and gain of accessory segments in these Gram-negative structures has been explained by the actions of IS*26*, and it appears that IS*1216* played a similar role in shaping the accessory content of pHHEf1 and pHHEf2.

### pRUM-like plasmids are internationally distributed and carry diverse accessory content

To assess the extent of structural variation in pRUM-like plasmid sequences, and to explore the role of IS*1216* in generating diversity, we screened the GenBank non-redundant nucleotide database with the pRUM replication initiation gene (last search: 23/11/2023). A total of 150 pRUM-like plasmids were identified and further characterised alongside pHHEf1 and pHHEf2 (Table S1). pRUM-like plasmids were found in genomes of *E. faecium* strains that were isolated between 1997 and 2022 from 20 countries on five continents, including the UK, USA, Japan, Egypt, Australia and India (Figure 2A). Sources of isolation included a variety of human clinical samples, dog faeces, and marine sediment. Host *E. faecium* belonged to a total of 25 different sequence types (STs), 24 of which were CC17. The most commonly represented STs were ST80, ST736, and ST17 (Figure 2C). The only non-CC17 type represented in the set was an ST266 strain that was isolated from dog faeces in The Netherlands.

**Figure 2:**
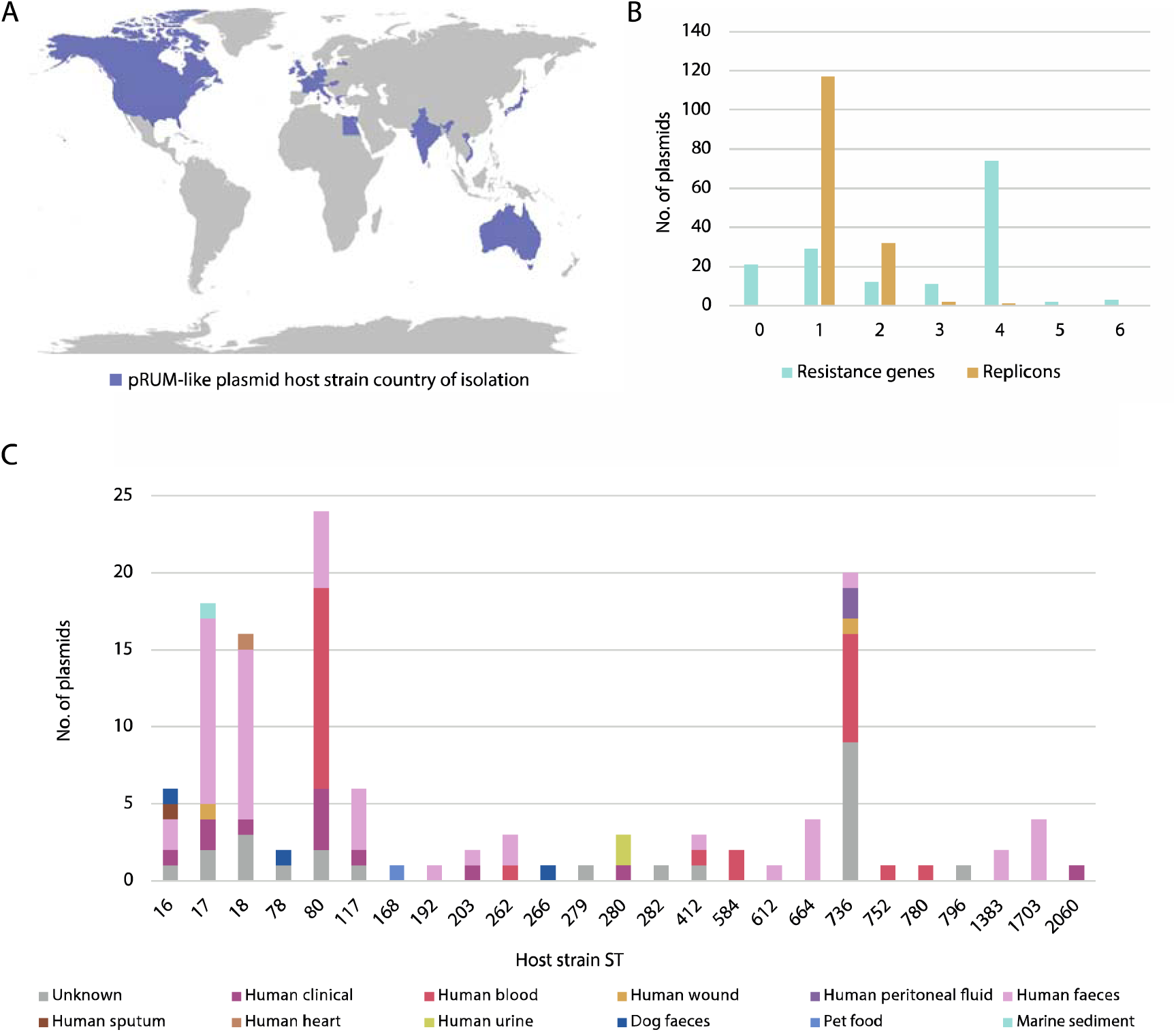
Overview of pRUM-like plasmid characteristics. A) Map displaying the countries of isolation for pRUM-like plasmid hosts examined here, shaded in purple. B) The number of antibiotic resistance genes (blue) and plasmid replicons (yellow) detected in pRUM-like plasmids using ResFinder and PlasmidFinder, respectively. The number of replicons includes the rep17 (pRUM) replicon that was detected in all 152 plasmids. C) The sequence type (ST) and isolation source of pRUM-like plasmid hosts. Data visualised in this figure can be found in Table S1.

Screening with ResFinder revealed 131 of the 152 pRUM-like plasmids contained one or more ARGs, with a range of 1-6 ARGs per plasmid (mode=4; 74 plasmids). These ARGs included all of those found in pHHEf1 and pHHEf2, as well as *vanB*, *aac(6’)-aph(2”)*, and tetracycline resistance genes *tet*(L), *tet*(M) and *tet*(S) (Table S1). As pHHEf2 is a cointegrate, we screened the collection for additional plasmid replicons using PlasmidFinder to detect further instances where pRUM-like plasmids have formed cointegrates with plasmids of recognisable types. Additional replicons were detected in 35 plasmids, with 32 containing one extra, two containing two, and one containing three (Figure 2B). The acquired replicons were of 10 different types (Table S1).

All pRUM-like plasmids contained at least one copy of IS*1216*. Screening for the boundaries of the accessory region revealed that those present in pRUM were maintained in just 16 plasmids, with IS*1216*-mediated deletions to the left or right responsible for their absence from the remainder. Two deletion events were found to the left of the accessory region, removing 108 bp or 319 bp of adjacent backbone, and at least 19 deletion events were identified to the right, removing between 35 bp and 3,578 bp of the backbone in that direction (Figure 3A). Some deletion events to the right of the accessory region have been large enough to remove the *axe*-*txe* genes, explaining reports of their variable presence. The backbone and copies of IS*1216* at the boundaries of the accessory region have been interrupted on multiple occasions by the insertion of nine different IS. Three IS were found in the accessory region-bounding copies of IS*1216*, and eight were found in the backbone (Figure 3A). In one plasmid (GenBank accession LR135359), the backbone has been interrupted by the insertion of Tn*1549*, which includes the *vanB* vancomycin resistance determinants. All other structure and content variation occurred within the accessory region.

**Figure 3:**
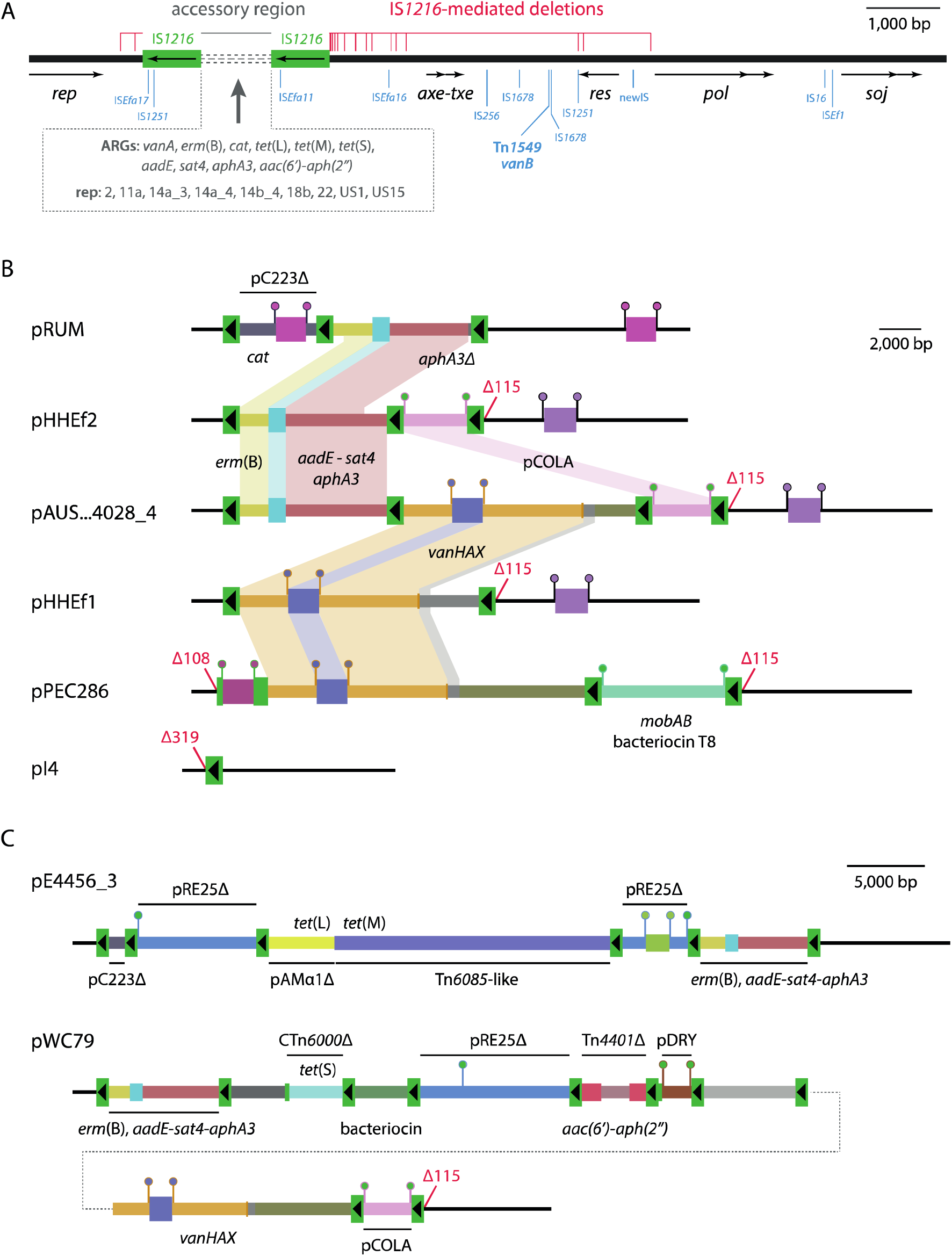
Variation in pRUM-like plasmids. A) Schematic summary of variation detected across pRUM-like plasmids in this study. The pRUM backbone is shown as a thick black line, with the extents and orientations of backbone genes indicated by labelled arrows below. The location of the accessory region is indicated above, with the IS*1216* at its boundaries shown, and the variable part of the region displayed as a set of grey dashed lines. Antibiotic resistance genes (ARGs) and plasmid replicons (rep) found in the accessory region are listed in the dashed box below. The extents of IS*1216*-mediated backbone deletion events are indicated by horizontal red lines above the backbone, with each vertical red line marking the extent of an individual deletion event. Insertions in the backbone or the IS*1216* at the boundaries of the accessory region are marked by blue lines, labelled with the identities of the elements involved. Backbone sequence drawn to scale from GenBank accession AF507977, with the positions of insertions and deletions determined from various plasmids listed in Table S1. B) and C) Scaled, linear maps of pRUM-like plasmids showing variation in accessory region content.

### IS*1216*-mediated cointegration has diversified pRUM-like plasmid accessory content

The first description of pRUM noted that it was a cointegrate, containing a segment related to the rolling-circle plasmid pC223 from *S. aureus*^11^. The pC223-like segment in pRUM is flanked by directly oriented copies of IS*1216* and contains the chloramphenicol resistance gene *cat* along with an origin-of-transfer (*oriT*), the mobilisation gene *mobC*, and a truncated *mobA* (Figure 3B). With IS*16* and one copy of its TSD removed, the 2,393 bp pC223-like fragment matches a contiguous segment of pC223 (GenBank accession AY355285) with nucleotide identity 98.9% (2366/2393). The absence of an internal TSD and part of the pC223 backbone is consistent with a 1,587 bp IS*1216*-mediated deletion event that occurred after the integration of the pC223-like plasmid.

Comparing the structures of pRUM and pHHEf2 revealed that they both include a pseudo-compound transposon (PCT) containing the *erm*(B) and *aadE*-*sat4*-*aphA3* genes, but that it differed due to internal deletions by IS*1216*, with the PCT in pRUM containing only a partial *aphA3* (Figure 3B). In contrast, the PCT in pHHEf2 was identical to that in pAUSMDU00004028_4, which also contains pCOLA and a variant of the Tn*1546*-derived *vanHAX* PCT carried by pHHEf1 (Figure 3B). The structures of the pHHEf1, pHHEf2 and pAUSMDU00004028_4 accessory regions are consistent with diversification by three mechanisms involving IS*1216*: acquiring TUs by the targeted conservative mechanism, acquiring a small plasmid by the copy-in mechanism, or losing TUs by homologous recombination between IS*1216* copies. The complete loss of cargo from the accessory region has resulted in the structure of strain E745 plasmid pI4, which is 10,763 bp and contains only a single copy of IS*1216* (Figure 3B). This structure could only have been generated by the loss of segments from an existing accessory region, as it has been demonstrated that IS*26* family elements do not move alone.^31^

Another example of a small plasmid capture event was found in pPEC286. The accessory region in pPEC286 includes an IS*1216*-bounded 6,181 bp segment that contains an internal 8 bp TSD consistent with copy-in cointegration (Figure 3B). With one copy of the TSD removed, the segment is identical to the complete sequence of the circular 6,173 bp plasmid pUCH1-4 (GenBank accession CP096215). Along with genes for replication (type rep11a) and mobilisation (*mobAB*), pUCH1-4 contains the gene for bacteriocin T8 and its cognate immunity gene (Figure 3B). Recent work has shown that bacteriocin T8 production by *E. faecium* provides a competitive advantage *in vitro* and in the mouse gut, and that bacteriocin T8 determinants are associated with emergent clinically relevant lineages globally.^32^ Only one further pRUM-like plasmid in the set examined here contained the same rep11a replicon (OR298096), but an IS*1216*-mediated deletion event removed its copy of the bacteriocin T8 gene, leaving only its immunity gene (Table S1). Another plasmid (OP378678) contained the bacteriocin T8 and immunity genes but had lost the rep11a replicon in a different IS*1216*-mediated deletion event (Table S1).

The *vanHAX* (114/152 plasmids), *erm*(B) (101/152) and *aadE*-*sat4*-*aphA3* (92/152) genes located in IS*1216* PCTs noted above were the most common ARGs in the collection of pRUM-like plasmids studied here. Examining the contexts of rarer ARGs (≤5 plasmids each) revealed that they too had been acquired in IS*1216*-mediated events. In pE4566_3 (GenBank accession LR135484), an internal 8 bp TSD indicates that a segment of the plasmid pRE25 (rep2) has been integrated via the IS*1216* copy-in mechanism (Figure 3C). The pRE25-derived sequence in pE4566_3 is separated by an IS*1216*-bounded segment that contains the *tet*(L) and *tet*(M) tetracycline resistance genes. The tetracycline resistance segment is comprised of sequences with two distinct derivations – the *tet*(L) gene is part of a segment derived from the *E. faecalis* plasmid pAMα1 (GenBank accession AF503772),^33^ while the *tet*(M) gene is in a partial Tn*6085*-like element, relatives of which have been associated with the dissemination of *tet*(M) in *E. faecium*.^34^ The structure of this segment suggests that IS*1216* captured these tetracycline resistance genes together after the Tn*6085*-like element inserted in a pAMα1-like plasmid.

Three pRUM-like plasmids in this collection were found in isolates of ST5 *S. aureus* (Table S1). These plasmids, called pWC79, are almost identical to one another and have previously been the subject of a study that characterised the emergence of vancomycin and methicillin resistant *S. aureus* (VMRSA) in a USA health clinic.^35^ Examining the structure of pWC79 in the context of this study revealed that its accessory region is comprised of segments derived from multiple elements that have been assembled by IS*1216* (Figure 3C). pWC79 includes a segment of pRE25, but this was acquired in a distinct event to that which introduced pRE25 to pE4566_3, as the insertion target site in pE4566_3 is uninterrupted in pWC79. In addition to pCOLA, pWC79 contains another small plasmid, pDRY, which features an internal 8 bp TSD and was also acquired in a copy-in cointegration event. The *tet*(S) gene in pWC79 is located in an IS*1216*-bounded segment containing sequence derived from CTn*6000*, and the *aac(6’)-aph(2”)* gene is part of a truncated Tn*4001* that has been captured by IS*1216*.

### 1997 clinical isolate *E. faecium* E1162 harbours an ancestor of pRUM-like plasmids

The collection of plasmid sequences examined here provides evidence for extensive IS*1216*-mediated variation in the pRUM-like accessory region. However, although the accessory region is bounded by directly oriented copies of IS*1216*, the flanking plasmid backbone does not feature a TSD. This suggests that the acquisition of the accessory region was associated with an IS*1216*-mediated deletion event that removed part of the original backbone sequence along with one or both copies of the TSD. A plasmid ancestral to the pRUM-like lineage would therefore be expected to lack the IS*1216*-bounded accessory region, and to feature a larger backbone sequence, part of which is identical or near-identical to the pRUM-like backbone.

*E. faecium* E1162 is a vancomycin-sensitive ST17 strain that was isolated from a bloodstream infection in France in 1997.^36^ When the E1162 genome was screened for plasmid replicons, a pRUM-like *rep* gene was found in a circular 57,031 bp plasmid. This plasmid, pCANE, contained the entire 10.2 kb pRUM-like backbone, which was 99.85% identical (10259/10274 bp) to that of pRUM (Figure 4). Screening with ISFinder revealed that the pRUM-like backbone segment in pCANE was interrupted by a single copy of IS*Efa4*, which was found downstream of the *axe*-*txe* genes. pCANE contained an additional 44,866 bp region that was located in precisely the same position as the pRUM-like accessory region (Figure 4). No sequence identical or near-identical to this 44.9 kb segment was found in GenBank (last search 06/06/2025). The closest GenBank matches were to other *E. faecium* plasmids, which were 98.56% identical (CP027503 and CP027514; 94% coverage). Two identical copies of a novel IS*110* family element, here named IS*Efa20*, were the only IS found in the 44.9 kb segment. Notably, IS*1216* was not present in pCANE. Additional content in pCANE outside the pRUM-like backbone was not associated with transposase genes or any recognisable MGE, and might therefore represent the proposed ancestral backbone sequence. We found 32 open reading frames in the 44.9 kb segment, many of which were putative transfer determinants. These included genes for pilus/fimbrae formation (pilin, isopeptide and sortases), DNA processing and transfer (a relaxase, coupling protein and ATPase) and establishment in recipient cells (anti-restriction and ssDNA-binding).

**Figure 4:**
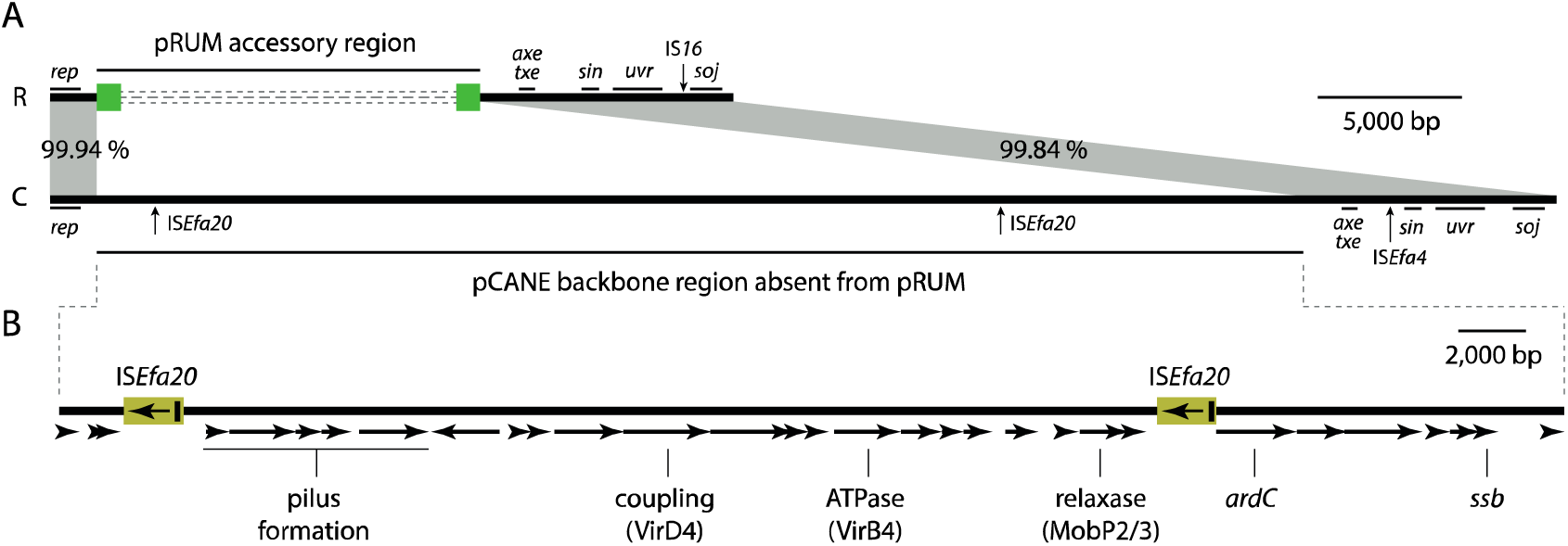
Ancestral plasmid pCANE. A) Scaled, linear maps of pRUM (R) and pCANE (C). The pRUM accessory region is shown as green boxes representing IS*1216* flanking dashed grey lines that represent variable segments. In both plasmids, the positions of backbone insertions are indicated by vertical arrows labelled with the name of the inserted element. The positions of regions of the pRUM backbone with known functions are marked with labelled vertical lines. Grey shading between the plasmids represents shared sequences, with their nucleotide identities (%) shown. B) pCANE backbone segment absent from pRUM. Open reading frames are shown as horizontal arrows, with predicted functions labelled.

## Discussion

Here we have characterised structural variation associated with diversification in the pRUM-like lineage of MDR plasmids, found primarily in *E. faecium*. pRUM-like plasmids have been associated with antibiotic resistance in the species for over two decades, but this is the first study to investigate their diverse accessory content from a topological perspective, and to trace the evolution of the lineage to a putative ancestor. We have outlined the crucial role of the IS*26* family element IS*1216* in the emergence and evolution of this lineage, mediating the loss of an ancestral transfer region, as well as the acquisition and continuing modulation of accessory segments. This has provided insights into the emergence of an important MDR platform, and more widely into the role of plasmid lineages in the success of bacterial clones that harbour them.

pRUM-like plasmids are comprised of an approximately 10 kb backbone and an IS*1216*-bounded accessory region that ranged from 809 bp (a single IS*1216*) to 285 kb in the collection of 152 plasmids examined here (Figure 3, Table S1). The pRUM-like backbone has been interrupted by the insertion of multiple IS, and the copies of IS*1216* at the boundaries of the accessory region have deleted adjacent backbone segments on multiple occasions (Figure 3A). However, the essentiality of some backbone genes to replication and stable maintenance has constrained the locations and extents of insertion and deletion events. Apart from the *vanB* determinants that have been acquired once through insertion of Tn*1549* in a backbone position, all other functional accessory genes have been acquired in the IS*1216*-bounded accessory region (Figure 3). The accessory region and its associated IS*1216* are therefore critical for the extensive diversification of the pRUM-like lineage.

We conclude that pCANE, carried by an ST17 *E. faecium* clinical isolate from 1997, is an ancestor of pRUM-like plasmids. pCANE does not contain the pRUM-like IS*1216*-bounded accessory region, but instead contains a unique 44.9 kb backbone segment (Figure 4). This segment includes putative transfer determinants, which suggests that the ancestors of pRUM-like plasmids were conjugative. The evolution of pRUM-like plasmids from a pCANE-like structure can be explained by events mediated by IS*1216* (Figure 5). First, a copy-in event would have introduced an IS*1216* TU to the pCANE backbone, generating a PCT structure flanked by an 8 bp TSD (Figure 5A). With the available data, it is not possible to determine the precise backbone location of this insertion, or whether one of the TSD copies remains in the sequence of pRUM. The loss of the 44.9 kb backbone segment, and one or both copies of the 8 bp TSD, likely occurred in one or more IS*1216*-mediated deletion events to generate a pRUM-like structure (Figure 5B). Further deletion events have continued to degrade the original backbone, as evidenced by the collection examined here (Figure 3A).

**Figure 5:**
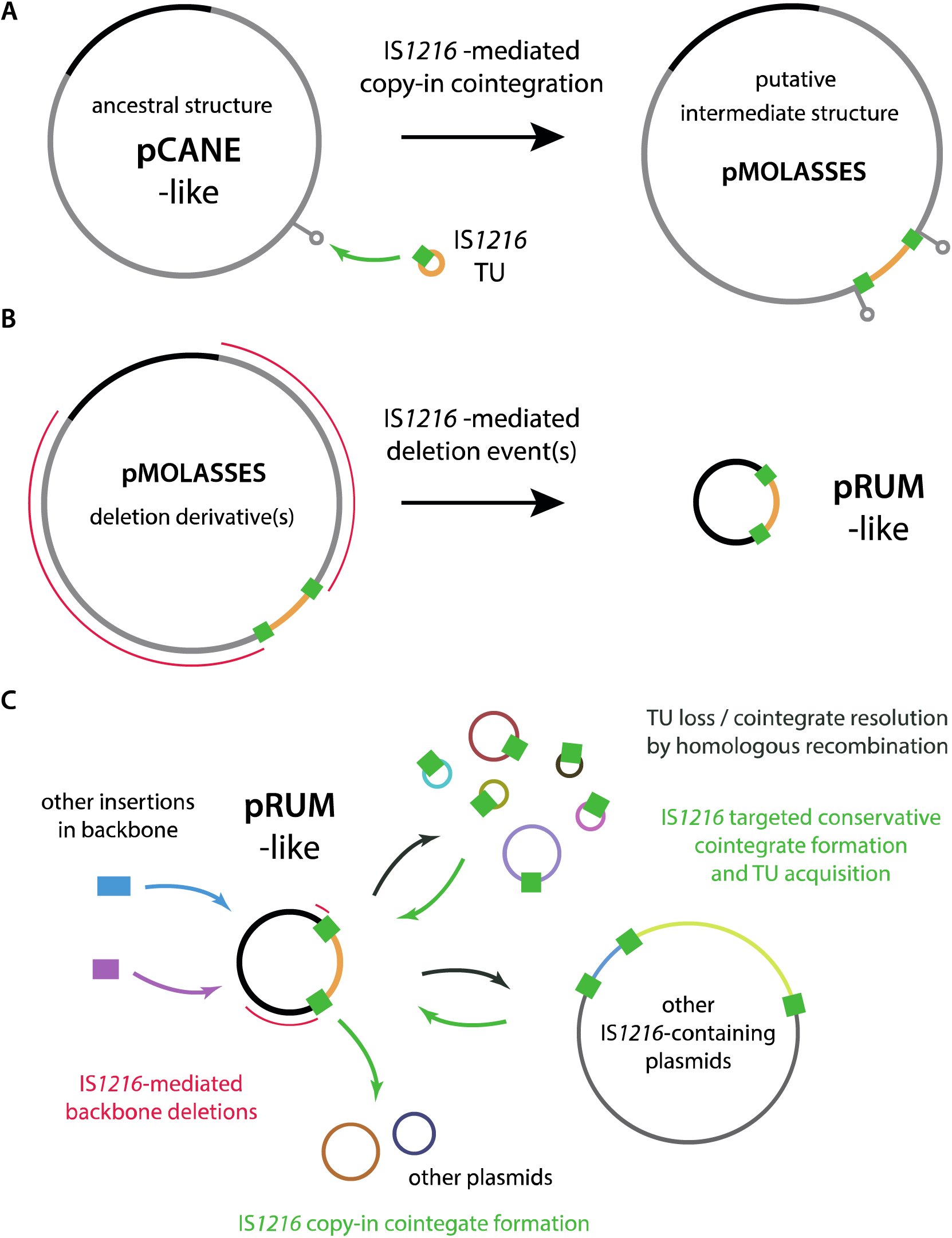
Emergence and ongoing evolution of the pRUM-like lineage. Proposed model for the evolution of pRUM-like plasmids from A) pCANE to the putative intermediate structure pMOLASSES, and B) from pMOLASSES to pRUM. Plasmids and a translocatable unit (TU) are shown as labelled circles with IS*1216* shown as a green box. A green arrow represents an IS*1216*-mediated copy-in cointegration event, and red lines represent IS*1216*-mediated deletion events. Elements are not drawn to scale. C) Overview of mechanisms contributing to the diversification of pRUM-like plasmids.

The loss of putative transfer determinants following the acquisition of an accessory region suggests that the emergence of the pRUM-like plasmid lineage involved an evolutionary trade-off, exchanging conjugative transfer ability for increased plasticity and access to a diverse pool of IS*1216*-associated genes. It is interesting to consider the consequences of this trade-off for the success of the pRUM-like lineage and its primary host, CC17 *E. faecium*. It is conceivable that the carriage of a stable and largely immobile MDR platform was an important contributor to the widespread success of CC17. In sweeping waves of selective pressure, as might be experienced in the human gut under courses of antibiotics, the pRUM-like plasmid-bearing CC17 would have an advantage over competing clones, while its MDR platform would be unlikely to transfer to those competitors. The absence of a transfer region in pRUM-like plasmids might provide another advantage to the host by reducing the risk of bacteriophage predation in the gut. Conjugative pili serve as extracellular targets for some bacteriophage, with plasmid carriage sensitising their hosts to infection.^37^ This risk would have been ameliorated by the loss of the pCANE genes that appear to determine a conjugative pilus (Figure 4). Despite these host advantages, horizontal mobility remains an important element in plasmid success, and re-establishing a degree of transfer capacity would be selfishly advantageous for the pRUM-like lineage. We have identified multiple cointegration events that have introduced putative conjugation or mobilisation determinants, and hypothesise that these acquisitions represent possible avenues for horizontal dissemination. These observations are similar to those made in *S. aureus*, where it has been demonstrated that the integration of small plasmids carrying *oriT* mimics is a common mobilisation strategy for larger non-conjugative plasmids.^8, 38^ Supporting the role of cointegrates in HGT, there is experimental evidence documenting the IS*1216*-mediated formation of a cointegrate comprised of pRUM-like and conjugative pHTß-like plasmids, facilitating transfer of the pRUM-like plasmid from *E. faecium* to *E. faecalis*.^39^

Beyond the pRUM-like lineage, our findings highlight the important role of IS*1216* in the shuttling of genetic material between *Enterococcus* and *Staphylococcus* species. Some ARG-containing passenger segments of IS*1216* PCTs within the pRUM-like accessory region are derived from distinct MGEs that were originally described in *Staphylococcus*, including the rolling-circle plasmid pC223 that carried *cat*, and the composite transposon Tn*5405* that carried *aadE*-*sat4*-*aphA3*.^40, 41^ The Tn*5405*-derived sequence in pRUM-like plasmids is part of a PCT that also includes the *erm*(B) gene (Figure 1), and this more complex structure was first found in veterinary isolates of *Staphylococcus intermedius*.^42^ In the opposite direction, IS*1216* was involved in the generation of a cointegrate pRUM-like plasmid that transferred from *E. faecium* to *S. aureus*, carrying the Tn*1546*-derived PCT and rendering the methicillin-resistant *S. aureus* recipient resistant to vancomycin (VMRSA).^35^ Clearly, cointegrate pRUM-like plasmids can function as natural shuttle vectors between these important nosocomial pathogens. Even when present only transiently in new hosts, pRUM-like plasmids can ‘shed’ IS*1216* TUs capable of inserting stably in new DNA molecules, or readily acquire new TUs from other molecules given the efficiency and self-preferential nature of the IS*26* family targeted conservative mechanism.^17^ pRUM-like plasmids therefore have concerning potential for the recruitment of further resistance genes, including the mobile linezolid resistance gene *cfr*(D) that has already been found associated with IS*1216* in clinical *E. faecium* isolates.^43, 44^

### Conclusions

The pRUM-like plasmid lineage is a stably maintained but highly plastic accessory gene platform in pandemic *E. faecium* CC17. This plasmid lineage appears to have emerged in ST17 in the 1990s, having been generated by the insertion of an IS*1216*-bounded accessory region. Subsequent IS*1216*-mediated events removed part of the ancestral plasmid backbone, including its putative transfer region. IS*1216* continues to drive diversification in pRUM-like plasmids by facilitating the acquisition of new DNA segments, most notably leading to the accumulation of ARGs. Cointegrate formation and the acquisition of horizontal transfer determinants from other plasmids represents a potential avenue for the dissemination of pRUM-like plasmids or their cargo into different clones or species.

## Data Availability

Whole genome sequencing reads for E1162 are available under NCBI BioProject PRJNA1279983. The complete sequences of plasmids pHHEf1, pHHEf2, and pCANE are available under GenBank accessions PV751131, PV763145, and CP195299, respectively.

## Author contributions

The project was conceived by RAM. E1162 was provided by WvS. Long-read sequencing was carried out by FA. Sequence annotation, database screening and metadata collection was performed by FA and RAM. Bioinformatic analyses were carried out by RAM, FA, and RSM. Data visualisation was by RAM and FA. All authors engaged in discussion of the results over the course of the study. The manuscript was drafted by RAM and FA, and was edited and revised by all authors.

## Acknowledgements

Thanks to Dr Abid Hussein for providing strains OI5 and OI17, and to Dr Lisa Lamberte for assistance with sequencing library preparation.

